# Sex-specific changes in gene expression and delayed sex differentiation in response to estrogen pollution in grayling (Salmonidae)

**DOI:** 10.1101/135210

**Authors:** Oliver M. Selmoni, Diane Maitre, Julien Roux, Laetitia G. E. Wilkins, Lucas Marques da Cunha, Etienne L. M. Vermeirssen, Susanne Knörr, Marc Robinson-Rechavi, Claus Wedekind

## Abstract

The synthetic 17α-ethinylestradiol (EE2) is an estrogenic compound of oral contraceptives and therefore a common pollutant that has been suspected to affect the demography of river-dwelling salmonids. We study a population of European grayling (*Thymallus thymallus*) that suffers from sex ratio distortions. Here we test how ecologically relevant concentrations of EE2 affect sex-specific gene expression around early stages of sex differentiation. We collected gametes from F1s of wild spawners, used them for *in vitro* fertilizations, and raised the resulting embryos singly under experimentally controlled conditions. Embryos were either exposed to 1ng/L EE2 or sham-exposed. RNA was collected from samples taken 10 days before hatching, at the day of hatching, and towards the end of the yolk-sac stage, to study gene expression and relate it to genetic sex (sdY genotype). We found that EE2 affects gene expression of a very large number of genes especially at the day of hatching. The effects of EE2 on gene expression is strongly sex-specific. At the day of hatching, EE2 affected about twice as many genes in females than in males, and towards the end of the yolk-sac larval stage, EE2 effects were nearly exclusively observed in females. Among the many effects was, for example, a surprising EE2-induced molecular masculinization in the females’ heads. Histological examination of gonadal development of EE2-treated or sham-exposed juveniles during the first 4.5 months after hatching revealed a delaying effect of EE2 on sex differentiation. Because grayling sex determination goes through an all-male stage (a rare case of undifferentiated gonochorism), the rate of EE2-induced sex reversal could not be unequivocally determined during the observational period. However, two EE2-treated genetic males had ovarian tissues at the end of the study. We conclude that common levels of EE2 pollution affect grayling from very early stages on by interfering with male and female gene expression around the onset of sex differentiation, by delaying sex differentiation, and by feminizing some males.

**Author contribution:** MRR and CW initiated the project. OS, DM, LW, LMC, and CW sampled the adult fish, did the experimental *in vitro* fertilizations, and prepared the embryos for experimental rearing in the laboratory. All further manipulations on the embryos and the larvae were done by OS, DM, LW, and LMC. The RNA-seq data were analyzed by OS, JR, and MRR, the histological analyses were done by DM, supervised by SK, the molecular genetic sexing was performed by OS and DM, and EV supervised the EE2 analytics. OS and CW performed the remaining statistical analyses and wrote the first version of the manuscript that was then critically revised by all other authors.

## 1. Introduction

Endocrine-disrupting chemicals are common pollutants that typically enter the environment after wastewater treatment. One of the most potent of these micropollutants is the synthetic 17-alpha-ethinylestradiol (EE2) that is used in oral contraceptives and hormone replacement therapies, and that is more stable and persistent than the natural estrogen it mimics [1]. EE2 concentrations of 1ng/L and higher have been found in river or lake surface waters [2] in lake sediments [3], and even in groundwater [4].

Exposure to 1 or few ng/L EE2 can be damaging to fish at various developmental stages. Embryos and early larvae can suffer from increased mortality, reduced growth, or malformations when exposed to EE2 [5, 6]. In juveniles and adults, exposure to EE2 can affect the response to infection [7], increase the susceptibility to other micropollutants [8], generally reduces growth and fertility [8, 9], and can even induce transgenerational effects on behavior and fertility in F1 [10] and F2 progeny [11]. Studies with experimental populations kept in 1,100 L ponds revealed population declines at concentrations of 1ng/L EE2 [12], and long-term, whole-lake experiments revealed significant ecosystem changes after experimental addition of 5-6 ng/L EE2: local populations of small fish declined (one species nearly got extinct), average body conditions of other fish, including top predators, declined significantly, and the prevalence of some zooplankton and insect species seemed to increase [13].

Experimental exposure to EE2, for example of juvenile sticklebacks (*Gasterosteus aculeatus*) to 35-40 ng/L, or of juvenile coho salmon (*Oncorhynchus kisutch*) to 2 or 10 ng/L, are associated with significant down- and up-regulations of various physiological pathways [14, 15]. Some of these effects on gene expression may be linked to the toxic effects of EE2 observed in juveniles and adults. However, it is likely that EE2 effects on gene expression depend on life history and on the developmental stage of an individual, i.e. on the timing of some physiological pathways in the organism. One important physiological pathway in this context is sex determination and gonad formation.

Sex determination is probably best seen as a threshold trait, with processes that occur early in development determining later processes [16]. In amphibians and fishes, these early processes can be very labile, i.e. potentially modifiable by external factors, even if they often have a clear genetic basis [17, 18]. Among these external factors that interfere with these early steps of sex determination are temperature or endocrine disrupting chemicals such as EE2 [17, 19]. When applied at the beginning of sex determination, they can tip the balance and cause a sex reversal, i.e. a mismatch between genotypic and phenotypic sex. Such EE2-induced sex reversals are sometimes but not always observed. If they are observed, the timing of exposure seems more relevant than the concentration: Orn et al. [20] found, for example, nearly complete sex reversal in zebrafish exposure to 5 ng EE2/L during embryo and larval stages, while exposure to 5-20 ng EE2/L at later stages did not seem to induce sex reversal in the same species [21].

Apart from gonad development, there can be many other fundamental differences between male and female development. Males and females may differ, for example, in average growth, timing of maturation, habitat use, or susceptibility to various stressors including infections [22]. It is therefore possible that effects of EE2 on the organism crucially depend on whether or not the organism is exposed to EE2 during the early steps of the sex determination cascade, when sex reversal is still possible [23].

Exposure to EE2 has been found to induce sex-specific effects in juvenile and adult fish [23]. These sex-specific effects can be large, as expected by the significant differences in male and female physiology. It remains unclear whether large differences should also be expected between the genetic sexes if their gonadal development may have been influenced by the sex-reversing effects of EE2 [24, 25]. Such questions can be studied if reliable sex-linked genetic markers are available for a given study species.

Here we study the sex-specific effects of ecologically relevant concentrations of EE2 on gene expression and gonad development in grayling (*Thymallus thymallus*), a river-dwelling salmonid fish that may often be exposed to EE2 pollution. Yano et al. [26] established sex-linked genetic markers that could be used to determine the genetic sex of many salmonids, including their sample of grayling taken from a fish farm in France. These markers could be successfully verified in over 100 phenotypically sexed adult grayling sampled from our study population [27]. We therefore use them here to separate effects of EE2 on gene expression in genetic males and females. Maitre *et al*. [27] found large effects of genetic sex on gene expression around the time of hatching from eggs, while gene expression did not seem to differ significantly at late embryogenesis. Their findings thus suggest that the physiological cascade of sex determination starts during embryogenesis and before hatching. We therefore study the interaction between EE2 and genetic sex on gene expression in embryos and larvae. Within-family comparisons are used to minimize potential effects of genetic variation. Possible interactions between EE2 and genetic sex on gonad development are studied histologically on samples taken over a period of several months.

## 2. Methods

### 2.1 Experimental breeding and raising

Mature males and females were sampled from a captive breeding stock and stripped for their gametes. These fish are F1 of the natural population described in Wedekind et al. [28]. Their gametes were used in two full-factorial breeding blocks. For each breeding block, 4 females were crossed *in vitro* with 5 males, i.e. 40 (2x4x5) different sibgroups were produced. After egg hardening for 2 hours, the fertilized eggs were transported to the laboratory where they were washed and distributed singly to wells of 24-well plates (Falcon, Becton-Dickinson), following the methods of von Siebenthal et al. [29]. The wells had been filled with 1.8 mL chemically standardized water [30] that had been oxygenated and temperated before use.

The embryos were incubated at 7°C and left undisturbed until 14 days post fertilization (dpf), when embryos were exposed either to 1 ng/L EE2 (by adding 0.2 mL water with a concentration of 10 ng/L EE2, see Brazzola *et al*. [6] for details), a strain of *Pseudomonas fluorescens* (“PF”; 10^6^ bacterial cells per well, the low-virulence strain PF1 in Clark et al. [31]), simultaneously to EE2 and *P. fluorescens* (EPF), or sham-treated (“control”, i.e. only adding 0.2 mL standardized water). A sample of 250 live embryos (EE2-treated or sham-treated offspring of one haphazardly chosen female that had been crossed with five males) was separated for analyses of gene expression, such that we had 25 embryos for each of the five sibgroups and the two treatments “EE2” and “controls”. The remaining embryos were raised until 40 dpf, i.e. until several days after hatching, when a subset of each treatment group was distributed to two 200 L tanks each filled with lake water (pumped from Lake Geneva at 40 m depth), fed with live *Artemia* and copepods and later dry food, and sampled in 5 monthly intervals for histological examination of the gonads (see Maitre *et al*. [27] for a description of sex differentiation). For the EE2-treated groups, 200 ng EE2 were dissolved in 200 L tanks each to reach a starting concentration of 1 ng/L. Every 7 days from then on, 40 L (i.e. 20%) were replaced with fresh lake water spiked with 40 mL of a 1 μg/L EE2 stock solution (i.e. 40 L at 1 ng/L EE2). Water samples (100 mL each) were then taken from each of the 4 EE2-treated tanks 1 hour after this weekly water exchange (T_0_) and 7 days later, just before the next water exchange (T_7_). These water samples were immediately frozen and stored at -20°C protected from light. Four consecutive T_0_ and 4 consecutive T_7_ samples were each pooled per tank for later determination of EE2 concentrations, i.e. EE2 concentrations were determined for the 4-week intervals these pooled samples covered, starting 47 dpf, 75 dpf, 103 dpf, and 131 dpf, respectively.

To quantify EE2, the water samples were thawed and filtered over glass fibre filters, their volume was set to 250 mL and the pH to 3. Four ng/L of EE2 D4 was added as internal standard and samples were enriched on LiChrolut EN / LiChrolut RP-C18 cartridges that had been conditioned with hexane, acetone, methanol and finally water (pH 3) [32]. After sample enrichment, cartridges were dried with nitrogen and eluted with acetone and methanol. Subsequently, solvents were changed to hexane/acetone 65:35 and extracts were passed over Chromabond Silica columns [33] and set to a volume of 0.25 mL. LC-MS/MS analysis was performed on an Agilent6495 Triple Quadrupole. An XBridge BEH C18 XP Column, 2.5 μm, mm X 75 mm and an acetonitrile / water gradient was used for liquid chromatography followed with post-column addition of ammonium fluoride solution. EE2 was quantified by monitoring the mass transition of 295 to 269, the transition of 295 to 199 served as a qualifier (internal standard was quantified at the following transitions: 299 to 273 and 299 to 147).

EE2 concentrations in 24-well plates and uptake by embryos are described in Marques da Cunha et al. (unpublished manuscript). Briefly, plates with newly fertilized embryos were spiked with EE2 to a concentration of 1 ng/L of EE2 in the plate wells. Analogously, plates without embryos were spiked to the same concentration. EE2 concentrations in the 24-well plates with and without embryos were measured in 5 time points across embryo development (i.e. from 1 dpf until shortly before embryo hatching). While EE2 concentrations have remained around 1 ng/L in plates without embryos throughout the 5 time points (indicating no apparent degradation or sorption of EE2 in 24-well plates), in plates with embryos, concentrations fell to below detection limit (<0.05 ng/L) before the first half of developmental time (indicating rapid embryo uptake).

In the 200 L tanks, median EE2 concentration were 0.33 ng/L at T_0_ and 0.11 ng/L at T_7_, corresponding to a median reduction of 66% of the EE2 dissolved in water over 7 days (see Supplementary Figure S1). We found no significant effects of sampling period on the EE2 measures at T_0_ (ANOVA, F_3_ = 1.20, p = 0.35) nor on the weekly reduction of EE2 in the tanks (F_3_ = 1.88, p = 0.19; excluding an unexplained outlier, see Figure S1 for discussion). The median EE2 concentration across T_0_ and T_7_ samples in the EE2-treated tanks was 0.2 ng/L from the first water exchange on, i.e. from 47 dpf on (EE2 was also detected in samples from control aquaria, see Figure S1 for discussion).

For the gene expression analyses, the first sampling of 12 embryos per family and treatment took place at 21 dpf, i.e. well before hatching could be expected. Embryos were immediately transferred to RNAlater (Thermo Scientific, Reinach, Switzerland). At 27 and 28 dpf, the incubation temperature was raised to 10°C and 11.5°C, respectively, in order to induce and synchronize hatching. The second sampling took place at day of peak hatching, i.e. 31 dpf (8 embryos per family and treatment). The third sampling took place at 52 dpf (5 yolk-sac larvae per family and treatment). Hatchlings and yolk-sac larvae were narcotized with 0.5mL/L KoiMed (fishmed GmbH, Galmiz, CH) for five minutes and then decapitated. The heads were immediately transferred to RNAlater. All samples were stored at -80°C.

RNA was extracted using the QIAgen 96 RNeasy Universal Tissue Kit (QIAGEN, Hombrechtikon, Switzerland). Manufacturer instructions were followed except that centrifugation (Eppendorf 5804 R centrifuge with an A-2-DWP rotor; Eppendorf, Schönenbuch, Switzerland) was done twice as long at half the speed. Because the RNA extraction protocol did not include a DNase treatment, DNA traces inside the RNA samples were amplified to determine the sdY genotype [26] of each individual, using the 18S gene as PCR internal control, either in a multiplex reaction used for samples with high amount of DNA, or after a second PCR amplification in single reactions with half the amounts of the respective primers each for samples with low DNA content (see Maitre *et al*. [27] for a more detailed protocol). Based on the sdY genotype, one female and one male per family and treatment group was haphazardly chosen for further analyses (in 2 of in total 30 combinations of family x treatment x time point, two females or two males, respectively, were used because only one sex could be found in the sample).

The RNA extracts were provided for library preparation in a equimolar concentration of 6 ng/μL in 100 μL of volume. 50 μL (*i.e.* 300 ng of RNA) were each used for library preparation on a robot using the Truseq Stranded RNA protocol (Illumina, Part# 15026495 Rev. A), then introduced in the Illumina sequencing platform (HiSeq 2500) for 100 cycles of multiplexed paired-end reads sequencing. The total 60 samples were sequenced in ten lanes (six samples per lane).

### 2.2 Bioinformatics pipeline

The RNA-seq analysis steps are described in Maitre *et al*. [27]. To summarize, reads were quality trimmed or filtered, and PCR duplicates were removed, resulting in a set of 60 RNA libraries with, on average, 2*40 millions of 80 bp reads each (standard deviation of six million reads). Reads from 24 libraries were assembled using Trinity (version 2.0.3; [34]). The resulting assembly was filtered to remove sequences that did not have a significant match against the UniprotKB/Swiss-Prot database (version of October the 16th 2015) [35] using blastx (version 2.2.26; [36]). Reads from all libraries were pseudo-mapped onto the assembled transcriptome using Kallisto (version 0.42) [37], and the read counts of the different isoforms of each gene were summed up for principal component analysis (PCA) and differential expression analysis at the gene level. PCA was performed on TMM-normalized [38] log2(count-per-million) values (CPM). Differential expression analysis was performed using the limma-voom Bioconductor package (version 3.26.3) [39, 40], with samples quality weights [41], on CPM values that were additionally cyclic loess normalized. In the linear model we considered developmental stage, sex and treatment as a combined variable (with twelve possible levels) and sib-group as an independent variable. A linear model was then fit for each gene, coefficients and standard errors were computed for all the contrasts of interest. Q-values [42] were calculated for each gene, and a threshold of q = 0.15 was used to call differentially expressed genes. Transcripts were annotated by referring to the Gene Ontology (GO) terms (Biological Function) [43] associated to similar genes in zebrafish. An enrichment analysis of GO terms was performed on differentially expressed genes using the goseq Bioconductor package (version 1.22.0; [44])

RNA extraction quality, male sex associated locus PCR, RNA-sequencing reads quality, transcriptome assembly, gene expression principal component and differential gene expression within control individuals are shown in Maitre *et al*. [27].

## 3. Results

### 3.1 Differential gene expression

In order to test for sex-specific effects, we compared the changes in gene expression under EE2 treatment for individuals of the same sex at the same developmental stage (Table 1). Under EE2 treatment, at the embryo stage there was an altered expression of several hundred genes in males (Table 1a, Figures S2a, S3, S4, Table S1), but only of a few genes in females (Table 1a, Figure S2b). At hatching day males showed an alteration in the expression of nearly 12,000 genes (Table 1b, Figure S2c, S5, S6, Table S2), while females at the same time point displayed an alteration in the expression of over 24,000 genes (Table 1b, Figure S2d, S7, S8, Table S3). At the first feeding stage only very few genes appeared altered in males (Table 1c, Figure S2e), whereas in female still around 14,000 genes were affected (Table 1c, Figure S2f, S9, S10, Table S4).

**Table 1.** Number of genes that are differentially expressed (q<0.15) in males and females tested at (a) embryo stage, (b) hatchling stage, and (c) larval stage at the onset of exogenous feeding, each after they had been exposed to EE2 or sham exposed.

|  |  | sham-exposed<br>females | EE2-exposed<br>males | EE2-exposed<br>females |
| --- | --- | --- | --- | --- |
| a) Embryos |  |  |  |  |
|  | Sham-exposed males | 15<sup>1</sup> | 704 | 863 |
|  | Sham-exposed females |  | 3 | 32 |
| b) Hatchlings |  |  |  |  |
|  | Sham-exposed males | 25,372<sup>1</sup> | 11,619 | 2 |
|  | Sham-exposed females |  | 0 | 24,728 |
| c) Larvae |  |  |  |  |
|  | Sham-exposed males | 1,110<sup>1</sup> | 4 | 41 |
|  | Sham-exposed females |  | 5,533 | 14,395 |

<sup>1</sup> reported in Maitre *et al.* [27]

In Table 2, the sex-specific alterations in gene expression are split according to the direction of the changes. Around hatching, about 5,000 genes were up-regulated in EE2-treated males while down-regulated in EE2-treated females, and about 4,000 were down-regulated in EE2-treated males while up-regulated in EE2-treated females (Table 2). The remaining sex-specific reactions to the EE2 treatment were mainly up- or down-regulation in one sex while there was apparently no change in the other sex (Table 2). See Figures S11 and S12 and Table S5 for EE2 effects on gene expression in both, males and females.

**Table 2.** Number of genes that were upregulated, i.e. had a positive log fold change of expression with q > 0.15 (UP), experienced no significant change in expression (NO), or were downregulated (DO) under exposure to EE2.

|  |  | Males UP | Males NO | Males DO | Total |
| --- | --- | --- | --- | --- | --- |
| a) Embryos |  |  |  |  |  |
|  | Females UP | 0 | 12 | 0 | 12 |
|  | Females NO | 245 | 34,613 | 457 | 35,315 |
|  | Females DO | 1 | 18 | 1 | 20 |
|  | Total | 246 | 34,643 | 458 | 35,347 |
| b) Hatchlings |  |  |  |  |  |
|  | Females UP | 147 | 7,959 | 4,043 | 12,149 |
|  | Females NO | 1,341 | 8,267 | 1,009 | 10,617 |
|  | Females DO | 4,962 | 7,495 | 117 | 12,574 |
|  | Total | 6,450 | 23,721 | 5,169 | 35,340 |
| c) Larvae |  |  |  |  |  |
|  | Females UP | 3 | 8,589 | 0 | 8,592 |
|  | Females NO | 1 | 20,949 | 0 | 20,950 |
|  | Females DO | 0 | 5,803 | 0 | 5,803 |
|  | Total | 4 | 35,341 | 0 | 35,345 |

### 3.2 Does EE2 treatment feminize males and masculinize females?

After focusing on sex-specific gene expression changes induced by EE2 treatment, we compared control males against EE2-treated females and control females against EE2 treated males (Table 1). The aim of this analysis was to investigate whether the EE2 treatment would feminize males, masculinize females, or increase the differences in gene expression between sexes. At embryo stage, we found three genes differentially expressed between EE2 treated males and control females (Table 1a) and 863 genes between control males and EE2 treated females (Table 1a). On hatching day, we found no differences in gene expression levels between control females and males treated with EE2 (Table 1b) and only two genes differing between control males and EE2 treated females (Table 1b). At first feeding stage, EE2 treated males expressed 5,533 genes differently in comparison to control females (Table 1c), while gene expression in control males differed in 42 genes only from the gene expression of EE2-treated females (Table 1c). Overall, there does appear to be transcriptome evidence of feminization of males and of masculinization of females.

### 3.3 Gene ontology enrichment analysis

Tables S1-S5 show the top 25 Gene Ontology terms enriched among genes differentially expressed in the above-described contrasts. Figures S3-S12 illustrate the Gene Ontology terms significantly enriched, grouped by relatedness. Table 3 shows a summary take-home from these Gene Ontology enrichments. Interestingly, many differences affect developmental processes. In male embryos, we also notice response processes, whether endocrine, xenobiotic or immunological. Finally, in female larvae a wide range of processes are affected, including behaviour and digestion.

**Table 3.** Summary interpretation of the differential gene expression analysis. The characterization of the biological processes relies on the gene ontology enrichment analysis of differentially expressed genes. Feminization and masculinization represent the situation where few genes (<100) are detected as differentially expressed, under EE2 treatment, in comparison to control female or control male, respectively. See Supplementary Figures S2-S13 and Tables S1-S5 for more detailed information.

| Developmental stage | Sex differences in gene expression <sup>1</sup> | EE2-effects in males | EE2- effects in females |
| --- | --- | --- | --- |
| Embryos | Weak | On genes associated with central nervous system development, immune response development, endocrine system and response to xenobiotic stimuli. | Unclear. |
| Hatchlings | Strong | On genes associated with skeletal muscle function and locomotion, metabolism of fatty acids and carbohydrates, DNA-repair and hindbrain morphogenesis. Possible feminization. | On genes associated with immune system activation, DNA-repair, axonal growth and synaptic transmission. Possible masculinization. |
| Larvae | Strong | A few genes only. | On genes associated with DNA-repair, signal transduction, immune system activation, carbohydrate metabolism, mechanosensory behavior and digestion. Possible masculinization. |

<sup>1</sup> Results from Maitre *et al.* [27]

### 3.4 Sex differentiation

Exposure to EE2 delayed the onset of morphological sex differentiation while exposure to *P. fluorescens* (PF) showed no effects (Table 4). Figure 1 shows the rates of juveniles without any testis or ovarian tissues over the five monthly samples, separately for fish that were or were not exposed to EE2, to illustrate the delaying effects of EE2. First signs of morphological sex differentiation could be observed at the 2^nd^ sample, but still more than 30% of the fish showed no sign of morphological sex differentiation at the end of the observation period (Figure 1).

**Table 4.** Likelihood ratio test on rate of undifferentiated gonads explained by time point of sampling and exposure to ethinylestradiol (EE2), the bacterium *P. fluorescens* (PF), or both (EE2 x PF). N_total_ = 251.

| Effect | $\chi^2$ | d.f. | P |
| --- | --- | --- | --- |
| **Time** | **72.3** | **4** | **<0.0001** |
| EE2 | <0.1 | 1 | 1.0 |
| PF | <0.1 | 1 | 1.0 |
| **Time x EE2** | **9.5** | **4** | **0.05** |
| Time x PF | 0.9 | 4 | 0.92 |
| EE2 x PF | <0.1 | 1 | 1.0 |
| Time EE2 x PF | 4.7 | 4 | 0.32 |

**Figure 1.**
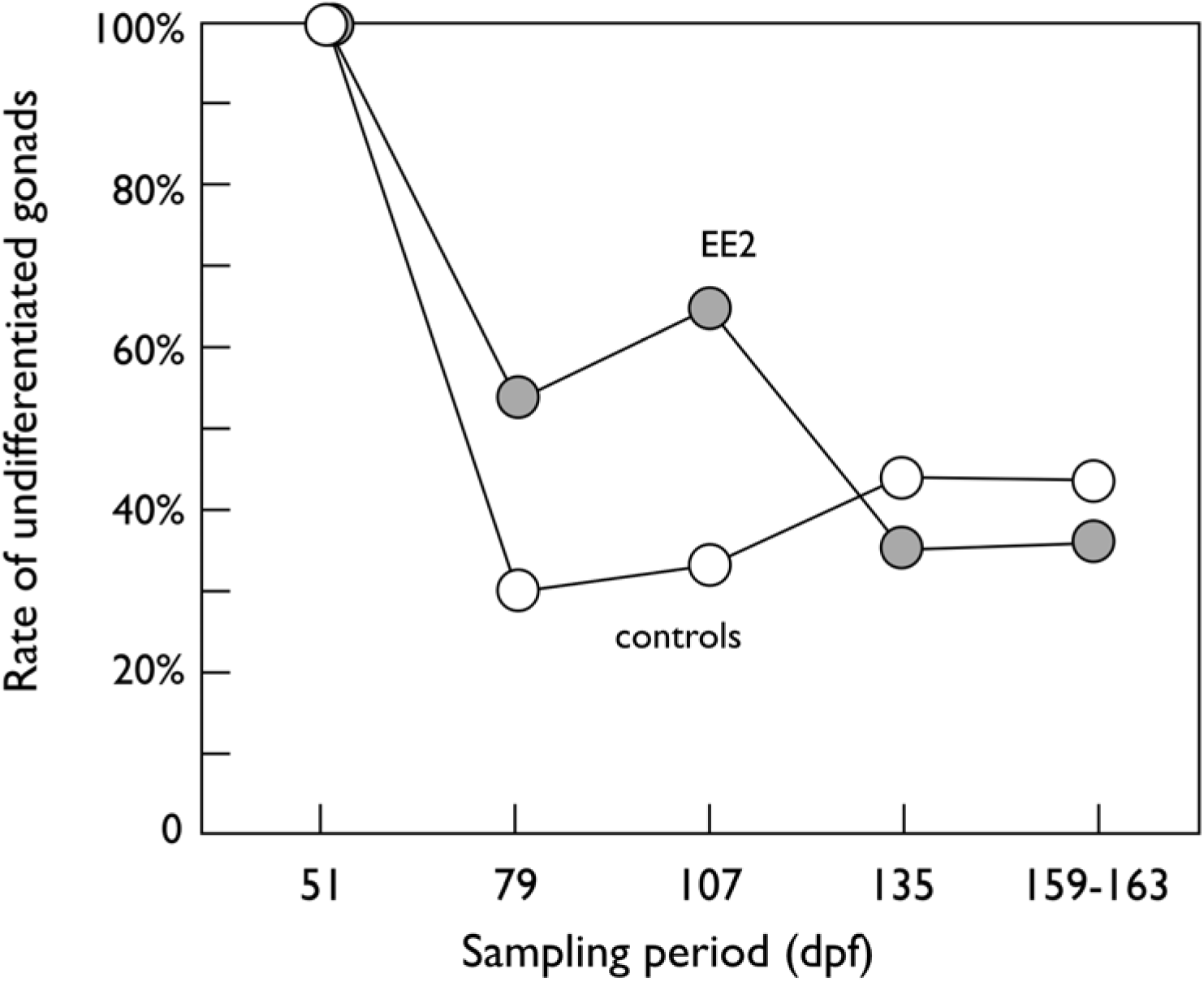
Rates of undifferentiated males and females (i.e. no testis nor ovarian tissues) when exposed to EE2 (closed symbols) or sham exposed (controls, open symbols), with or without additional exposure to *P. fluorescens* during embryogenesis each (exposure to the microbe showed no significant effects on sex differentiation). See text and Table 4 for statistics.

Among those fish where first stages of sex differentiation could be identified, only testis tissue could be observed at the 2^nd^ sampling (79 dpf), while the rate of ovarian tissue rose quickly to 70.8%, 75.9%, then 79.3% over the 3^rd^, 4^th^, and 5^th^ sampling periods, respectively. The rates of ovarian versus testis tissue did not differ between EE2-treated and sham-treated grayling (χ^2^ = 0.26, p = 0.61).

Genetic sexing of all 103 individuals of the 4^th^ and 5^th^ sample (135 dpf and 159-163 dpf) revealed a genetic sex ratio of 55.3% males that did not deviate from equal sex ratio (χ^2^= 1.2, d.f. = 1, p = 0.28). Equal sex ratios can therefore be assumed for all earlier samples. At these two last sampling days, all females except three showed ovarian tissue (ovaries or testis-to-ovaries). The three exceptions were from the sham-treated controls and showed testis tissue, i.e. no genetic female was undifferentiated at that these last sampling dates. In contrast, 45 of a total of 57 genetic males were still undifferentiated at that time, 10 showed testis tissue, one showed the testis-to-ovaries phenotype, and one had ovaries. The latter two males with ovarian tissues had both been EE2-treated.

## 4. Discussion

We tested and described the effects of exposure to low, ecologically relevant, concentrations of EE2 on sex-specific gene expression in embryos and larvae of grayling, a river-dwelling salmonid that is often exposed to this type of pollution. From what is known about possible EE2 effects on fish in general, we expected that this common micropollutant may (i) affect sex determination of grayling by influencing the few initial triggers that start the canalized developmental process of gonad formation, and (ii) be toxic to the embryos and larvae because it interferes with different types of physiological processes, especially those that are endocrinologically regulated (see references cited in the Introduction). We therefore expected EE2 to have significant effects on gene expression at various developmental levels, and we indeed found such effects at all the developmental stages we studied here. However, we had no clear *a priori* expectancy about whether EE2 would also affect the genetic males and genetic females differently at any of these stages.

We started from the premise that sex in gonochoristic species is a threshold trait, i.e. a canalized developmental process that has one or few initial triggers [16]. In grayling, the initial trigger (or triggers) that determine phenotypic sex happen during embryogenesis well before hatching, because over 25,000 genes are already differentially expressed between genetic males and females at the day of hatching [27]. The 15 genes that Maitre et al. [27] found to be differentially expressed in genetic males and females at the embryo stage 10 days before hatching suggest that sex determination starts around then, i.e. at a time when the embryos had already been exposed to EE2 for several days in the present study.

One possible scenario is hence that EE2 could tip the balance at the early steps of sex determination so that all individuals follow the developmental process that leads to the female phenotype regardless of their sdY genotype (i.e. sex reversal of genetic males). If so, EE2 would not be expected to show sex-genotype specific effects on gene expression during later stages of sex differentiation. However, we found strong interactions between genetic sex and EE2 on gene expression. These sex-specific reactions to EE2 also depended on the developmental stages we studied. At embryo stage, expression of only few genes seemed biased in genetic females, but gene expression in genetic males was already significantly affected, with about 700 genes up-or down-regulated under the influence of EE2. The outcome was somewhat reverse at the larval stage: now only few genes of genetic males seemed to be affected by EE2, while over 14,000 genes were differentially expressed in genetic females. An even more pronounced effect of EE2 could be seen at the day of hatching: far over 20,000 genes showed differential expression, and about 9,000 of them were either upregulated in genetic females and down-regulated in genetic males or down-regulated in genetic females and upregulated in genetic males.

The strong sex-specific responses to EE2 suggest that exposure to ecologically relevant concentrations of EE2 during embryogenesis did not simply tip the balance at early steps of sex determination so that all individuals would become phenotypic females and would show similar patterns of gene expression from then on. Instead, our observations suggest that genetic sex largely determined phenotypic sex, and that EE2 then interfered with sex-specific gene expression, creating the strong sex-specific reactions to EE2. This conclusion is supported by the observation that the low concentrations of EE2 we used are commonly observed in natural rivers and streams while there is little evidence for complete and population-wide sex reversal, even if natural populations sometimes show distorted sex ratios [45].

While the interaction between EE2 and genetic sex on gene expression suggested that exposure to EE2 is mostly interfering with the development of a phenotype that would correspond to the genotypic sex, we nevertheless found two genetic males that had developed ovarian tissues when exposed to EE2. It is possible that we missed some sex-reversal and the rate of sex reversal is higher, because we learned only during the course of the study that the grayling is a rare example (and probably even the only one so far) of an undifferentiated gonochorist that goes through an all-male stage before gonads differentiate into testes and ovaries [27]. Testis tissue in early juveniles can therefore not be interpreted as evidence for normal development of a male phenotype, and we probably stopped the study too early to get a reliable estimate of the rate of sex reversals in grayling exposed to EE2. However, by the end of the study, nearly all genetic females had developed ovarian tissue. This suggests that the rate of sex reversals is either low indeed, or that sex reversal would slow down gonad development so much that we would have missed many sex-reversed individuals within our observational window. We know of no examples or arguments in the literature that would support the latter possibility. However, we found that ecologically relevant concentrations of EE2 could affect sex determination in at least some individuals. It would therefore be interesting to learn what makes an early embryo susceptible or resistant against chemically induced sex reversal.

Our gene expression analysis suggested that exposure to EE2 induces effects in the transcriptomes of the brain that could be interpreted as partial sex reversal, especially around hatching time when many genes were differentially upregulated in genetic females and downregulated in genetic male, and vice versa, under the influence of EE2. At the day of hatching, genetic males appeared to have a somewhat feminized transcriptome when treated with EE2. This effect seemed to cease before the first feeding stage. Genetic females seemed to experience some form of masculinization of the transcriptome of the brain, at least from hatching day to first feeding stage.

Estrogens are known to affect functions of the nervous system, including synapsis homeostasis [46], neurogenesis [47], and sexual differentiation [47-49]. The latter association is particularly interesting here because estrogens have repeatedly been shown to induce, to some degree, a masculinization of the brain [47-49]. This is because androgenic hormones, produced mainly in the male gonads, are transformed by brain cells into estrogen that subsequently act locally via specific receptors to induce typical male development [48].

In salmonids, gonadal precursor cells typically differentiate during the weeks that follow hatching [17, 50]. During this period, the emergence of an endogenous synthesis of sexual hormones could explain why we observe a divergent response between sexes, especially if we consider that endocrine active compounds often elicits non-monotonic dose-responses [51, 52]. Such dose-effects could explain why we observed strong sex-specific responses to EE2 at hatching day and why these responses partly declined towards first feeding stage. Apart from the likely effects of EE2 on normal development of male and female phenotypes, exposure to EE2 also affected the expression of genes linked to many other physiological systems, including, for example, various aspects of the immune system, the endocrine system, the development of skeletal muscles, or of the digestive system.

In conclusion, exposure to ecologically relevant concentrations of EE2 during early embryogenesis may potentially tip the balance at early steps of sex determination so that all individuals follow the developmental process that leads to the female phenotype regardless of their sdY genotype (i.e. sex reversal of genetic males). If so, and if gene expression is then mainly determined by this gonad development, EE2 would probably not be expected to show strong sex-genotype specific effects on gene expression during later stages of sex differentiation. However, we found strong effects of both, EE2 and genetic sex, on gene expression at three different developmental stages shortly after the onset of sex determination. Such sex-specific effects of EE2 suggest that sex reversal has mostly not happened, although we found two genetic males with gonadal tissues at the end of the study. Our observations support earlier conclusions that ecologically relevant concentrations of EE2 act mostly toxic (i.e. affect growth and viability) but have little effect on sex determination in grayling [53], and that the observed sex ratio distortion may in the population [45] may be due to sex-specific survival instead rather than induced sex reversal.

## 5. Acknowledgements

We thank B. Bracher, U. Gutmann, C. Küng, R. Mani, M. Schmid, and T. Vuille from the Fishery Inspectorate Bern for permissions and provision of fish for experimental fertilizations, the *Pachtverein Thun* for catching the wild fish, the Saint-Sulpice pump station for zooplankton supplies, T. Braunbeck for access to the histology laboratory, the Genomic Technologies Facility of the University of Lausanne for library preparation and sequencing, and C. Berney, T. Bösch, J. Buser, E. Clark, B. des Monstiers, M. dos Santos, J. Kast, C. Luca, Y. Marendaz, D. Nusbaumer, D. Olbrich, E. Pereira Alvarez, M. Pompini, A.-L. Roulin, V. Sentchilo, R. Sermier, B. von Siebenthal, and D. Zeugin for assistance in the field or the laboratory. This project was authorized by the veterinary authorities of the cantons of Bern (BE118/14) and Vaud (VD2710; VD2956) and financially supported by the Swiss Federal Office for the Environment and the Swiss National Science Foundation (31003A_159579).

